# Cell fixation and preservation for droplet-based single-cell transcriptomics

**DOI:** 10.1101/099473

**Authors:** Jonathan Alles, Nikos Karaiskos, Samantha D. Praktiknjo, Stefanie Grosswendt, Philipp Wahle, Pierre-Louis Ruffault, Salah Ayoub, Luisa Schreyer, Anastasiya Boltengagen, Carmen Birchmeier, Robert Zinzen, Christine Kocks, Nikolaus Rajewsky

## Abstract

**Background:** Recent developments in droplet-based microfluidics allow the transcriptional profiling of thousands of individual cells, in a quantitative, highly parallel and cost-effective way. A critical, often limiting step is the preparation of cells in an unperturbed state, not compromised by stress or ageing. Another challenge are rare cells that need to be collected over several days, or samples prepared at different times or locations.

**Results:** Here, we used chemical fixation to overcome these problems. Methanol fixation allowed us to stabilize and preserve dissociated cells for weeks. By using mixtures of fixed human and mouse cells, we showed that individual transcriptomes could be confidently assigned to one of the two species. Single-cell gene expression from live and fixed samples correlated well with bulk mRNA-seq data. We then applied methanol fixation to transcriptionally profile primary single cells from dissociated complex tissues. Low RNA content cells from *Drosophila* embryos, as well as mouse hindbrain and cerebellum cells sorted by FACS, were successfully analysed after fixation, storage and single-cell droplet RNA-seq. We were able to identify diverse cell populations, including neuronal subtypes. As an additional resource, we provide ‘dropbead’, an R package for exploratory data analysis, visualization and filtering of Drop-seq data.

**Conclusions:** We expect that the availability of a simple cell fixation method will open up many new opportunities in diverse biological contexts to analyse transcriptional dynamics at single cell resolution.

## Background

A tissue is composed of many specialized cell types, each of which can have various biological states. Rather than studying global gene expression of a tissue as a whole, it has been recognized that transcriptional profiling at a single-cell resolution [1-4] provides a much more complete and accurate description of its biological function [5, 6]. Recent advances in droplet-based microfluidic technologies have made it possible to capture, index and sequence the transcriptional profiles of thousands of individual cells in a highly parallel, ultrafast and affordable manner [7, 8].

In the ‘Drop-seq’ method described by Macosko et al. [7], cells are separately encapsulated in nanoliter-sized droplets together with a single bead in a microfluidic device. One bead delivers barcoded primers, each harboring a polymerase chain reaction (PCR) handle, a cell barcode, and a multitude of different unique molecular identifiers (UMIs), followed by a polyT sequence. The beads are suspended in a lysis buffer, resulting in the cell being lysed upon droplet formation. Cellular mRNAs are released and can hybridize to the polyT sequences of the barcoded bead primers. After collection, the droplets are broken and the mRNA is reverse-transcribed into cDNA, PCR-amplified, and sequenced in bulk. Computational analysis allows to distinctly assign which mRNAs originate from the same cell by means of the cell barcode. The UMIs are used to identify and remove PCR duplicates and to digitally count distinct mRNA molecules.

Despite the rapid rise in high-throughput single-cell RNA-sequencing (RNA-seq) methods, including commercialized versions of automated platforms such as the Fluidigm C1, 10XGenomics or 1CellBiO systems, comparatively little attention has been given to the limitations that need to be overcome in the preparation and handling of cellular input material [9]. A major challenge in obtaining meaningful information is the use of a high-quality single-cell suspension, which appropriately reflects the transcriptional state of each cell within its natural or experimentally-intended environment. The steps between cell harvesting from culture or after tissue dissociation, isolation of single cells, and mRNA capture are particularly critical as they are prone to introduce transcriptome changes and degradation of RNA. Requirements such as the need to pool cells from several tissues or culture conditions, possibly combined with time course experiments, represent an additional restriction.

In principle, many of these problems could be addressed with the help of chemical fixation. Unlike aldehydes, methanol and ethanol are coagulating fixatives that do not chemically modify nucleic acids [10, 11]. Alcohols act by dehydration: in higher than 65% alcohol and in the presence of salts, nucleic acids occur in a collapsed state, that can be reverted to its original form by a simple rehydration. We have previously shown that fixation with 80% methanol is compatible with next-generation sequencing and library preparation for both, mRNAs and small RNAs [12]. Fixation was critically required for successful genome-wide gene expression profiling of FACS-sorted, one to four-cell stage *C. elegans* embryos, a complex tissue undergoing rapid and dynamic transcriptional changes [12].

Here, we adapted the methanol-based fixation protocol from Stoeckius et al. [12] to preserve cells for subsequent profiling of single-cell transcriptomes by Drop-seq. We first analyzed both live and fixed mixtures of cultured human (HEK) and mouse (3T3) cells to demonstrate that methanol fixation does not change the numbers of genes and transcripts (defined as the number of UMIs) detected per cell or interfere with unambiguous assignment of reads to one or the other species. We then applied methanol fixation to a larger scale analysis of ~ 9 000 primary cells from dissociated *Drosophila* embryos or sorted mouse hindbrain cells. We demonstrate that Drop-seq profiling of single-cell transcriptomes with methanol-fixed cells performs well with both cultured and primary cells.

Additionally, we provide a computational resource to facilitate the exploration of droplet-based single-cell sequencing data. ‘Dropbead’ can be readily used to visualize basic statistics and quantitative parameters, compare different samples and filter samples prior to subsequent analysis.

## Methods

### Preparation and fixation of cell lines for Drop-seq

Human Flp-In T-Rex 293 HEK cells were a gift from M. Landthaler (MDC, Berlin) originally obtained from Invitrogen (cat no. R78007); murine NIH-3T3 cells were from DSMZ (ACC 59). Cells were grown in DMEM (Invitrogen 61965-026) without antibiotics containing 10% fetal bovine serum, and confirmed to be mycoplasma-free (LookOut Mycoplasma PCR detection kit, Sigma Aldrich). Cells were grown to 30 to 60% confluence, dissociated with 0.05% bovine trypsin-EDTA (Invitrogen 25300062), quenched with growth medium, and further processed as described previously (Macosko et al. [7], Online-Dropseq-Protocol-v.3.1 http://mccarrolllab.com/dropseq/). Briefly, between ~ 1 to 10 × 10^6^ were handled always in the cold and kept on ice, pelleted at 300 × g for 5 minutes at 4°C, washed with 1× PBS + 0.01% bovine serum albumin fraction V (BSA) (100 µg/ml; Biomol 01400), resuspended in PBS, filtered through a 40 or 35 µm cell strainer and counted. For Drop-seq, a [1 + 1] mixture of [HEK + 3T3 cells] was prepared at a final (combined) input concentration of 100 cells/µl in 1× PBS + 0.01% BSA (corresponding to a final concentration of 50 cells/µl after mixing with lysis buffer in the co-flow device). Cells were trypsinized, and between 1 and 4 × 10^6^ cells were processed as described above for Drop-seq [7]. Cells were handled in regular (not: “low-binding”) microcentrifuge tubes to minimize cell loss, and kept cold at all times. After straining and counting, cells were pelleted at 300 × g for 5 minutes at 4°C, the supernatant was removed manually and the cell pellet resuspended in 2 volumes (200 µl) of ice-cold PBS.

Methanol fixation was adapted from Stoeckius et al. [12]. To avoid cell clumping, 8 volumes (800 µl) of methanol (grade p. a.; pre-chilled to -20°C) were added drop-wise, while gently mixing or vortexing the cell suspension. Methanol-fixed cells were kept on ice for a minimum of 15 minutes and were then stored at -80°C for up to several months, as indicated. For rehydration, cells were either kept on ice after fixation (Fixed) or moved from -80°C to 4°C (Fixed 1 or 3 weeks) and kept in the cold throughout the procedure. Cells were pelleted at 1000 to 3000 × g, resuspended in PBS + 0.01% BSA, centrifuged again, resuspended in PBS + 0.01% BSA, passed through a 40 or 35 µm cell strainer and counted, and diluted for Drop-seq in PBS + 0.01% BSA as described above. For control of RNA quality after fixation, cells were resuspended in PBS, kept on ice for 5 to 10 minutes, re-pelleted and then RNA was extracted with TRIZOL.

### Preparation of *Drosophila* cells for Drop-seq

The *D. melanogaster* strain used was *y ^1^ w^1118^; P{st.2::Gal4}; P{vnd::dsRED}* [13]. Eggs were collected on apple juice-agar plates for 2 hours and aged for ~6 hours at 25°C. Embryos were dechorionated for 1 min in ~4% NaOCl (diluted commercial bleach) and extensively washed with deionized water. Excess liquid was removed and embryos were transferred to 1ml ice cold dissociation buffer (cell culture grade PBS, 0.01% molecular biology grade BSA) 6 hours after embryo collection. Approximately 500 - 1000 embryos were collected prior to dissociation; a small subsample was stored in methanol for later staging by microscopy. Embryos were dissociated in a Dounce homogenizer (Wheaton #357544) with gentle, short strokes of the loose pestle on ice until all embryos were disrupted. The suspension was transferred into a 1.5 ml microfuge tube, cells were pelleted for 3 minutes at 1 000 g at 4°C. The supernatant was exchanged for 1 ml fresh dissociation buffer. Cells were further dissociated using 20 gentle passes through a 22G × 2” needle mounted on a 5 ml syringe. The cell suspension was then gently passed through a 20 µm cell strainer (Merck, NY2002500) into a fresh 1.5 ml microfuge tube, residual cells were washed from the strainer using a small amount of dissociation buffer. Cells were pelleted for 3 minutes at 1 000 g at 4°C and resuspended in 100µl fresh dissociation buffer and counted. Samples were fixed by adding 4 volumes of ice-cold 100% methanol (final concentration of 80% methanol in PBS), and thoroughly mixed with a micropipette. Cells were stored at -20°C until use (for up to two weeks).

For Drop-seq runs, cells were moved to 4°C and kept in the cold throughout the procedure. Cells were pelleted at 3 000g for 5 minutes, rehydrated in PBS + 0.01% BSA in the presence of RNAse inhibitor (RiboLock 1U/µl), pelleted and resuspended again in presence of RNAse inhibitor, passed through a 35 µm cell strainer, counted, and finally diluted for Drop-seq into PBS + 0.01% BSA (final concentration of 50 cells/µl).

### Preparation of mouse hindbrain cells for Drop-seq

Brains of newborn mouse pups (C57BL/6; postnatal day 5) were dissected in ice cold buffer (120 mM NaCl, 8 mM KCl, 1.26 mM CaCl_2_, 1.5 mM MgCl_2_, 21 mM NaHCO_3_, 0.58 mM Na_2_HPO_4_ and 30 mM Glucose, pH 7.4) that was saturated with 95% O_2_ and 5% CO_2_. Transsections at the level of the pons and C2 motor roots were performed using a razor blade to isolate the rhombencephalon. Hindbrain and cerebellar tissues were dissociated using the Papain Dissociation System (Worthington) according to the manufacturer’s instructions. To facilitate dissociation and prevent aggregation, DNAse I (Roche, 5U/ml) was added to the protease solution. After inactivation, cells were resuspended in Mg^2+^- and Ca^2+^-free Hank’s Balanced Salt Solution. Live (propidium iodide-negative) cells were sorted directly into ice-cold methanol (final concentration 80% methanol) and stored fixed cells for more than four weeks at -80°C. The sort was carried out under low pressure flow settings, previously optimized to maximize recovery of viable cells for subcultures. For Drop-seq, aliquots with 10^6^ or 3×10^5^ methanol-fixed sorted hindbrain cells were pelleted and processed as described above. RNAse inhibitor (RiboLock 1 U/µl) was added during the rehydration, wash and cell straining step. Cell recovery was 19% and 12% from the two cell preparations, respectively. Cells were diluted 1:3 into PBS-BSA 0.01% and then used for Drop-seq.

### Drop-seq procedure, single-cell library generation and sequencing

Monodisperse droplets of about 1 nl in size were generated using microfluidic polydimethylsiloxane (PDMS) co-flow devices (Drop-seq chips, FlowJEM, Toronto, Canada; rendered hydrophobic by pre-treatment with Aquapel). Barcoded microparticles (Barcoded Beads SeqB; ChemGenes Corp., Wilmington, MA) were prepared and, using a self-built Drop-seq set up, flowed in closely following the previously described instrument set up and procedures by Macosko et al. [7] (Online-Dropseq-Protocol-v.3.1 http://mccarrolllab.com/dropseq/). For most microfluidic co-flow devices, emulsions were checked by microscopic inspection; typically less than 5% of bead-occupied droplets contained more than a single barcoded bead. Droplets were collected in one 50 ml Falcon tube for a run time of 12.5 minutes. Droplets were broken promptly after collection and barcoded beads with captured transcriptomes were reverse transcribed without delay, then exonuclease-treated and further processed as described [7]. The 1^st^ strand cDNA was amplified (after assuming loss of about 50% of input beads) by equally distributing beads from one run to 24 or 48 PCR reactions (between 10 and 30 anticipated STAMPS per tube; 50 µl per PCR reaction; 4 + 9 cycles (except for mouse hindbrain replicate 2: 4 + 12 cycles). 20 or 10 µl fractions of each PCR reaction were pooled, then double-purified with 0.6x volume of Agencourt AMPure XP beads (Beckman Coulter, cat. no. A63881), and eluted in 12 µl H_2_O. 1 µl of the amplified cDNA libraries was evaluted and quantified on a BioAnalyzer High Sensitivity Chip (Agilent). 600 pg of each cDNA library was fragmented and amplified (12 cycles) for sequencing with the Nextera XT v2 DNA Library Preparation kit (Illumina) using custom primers that amplified only the 3′ ends [7]. Libraries were purified with 0.6x volume of AMPure XP beads, followed by 0.6× to 1× volume AMPure beads to completely remove primer dimers and achieve an average length ~500 to 700 bp, quantified and sequenced (paired end) on Illumina Nextseq500 sequencers (library concentration: 1.8 pM; 1% PhiX spike-in for run quality control; Nextseq 500/550 High Output v2 kit (75 cycles); read1: 20 bp (bases 1-12 cell barcode, bases 13-20 UMI) (custom primer 1 Read1CustSeqB), index read: 8 bp, read 2 (paired end): 64 bp).

Unique identifiers: Live: GSM2359902; Fixed: GSM2359903; Fixed 1 week: GSM2359904; Fixed 3 weeks: GSM2359905; *Drosophila*: mel_rep1: GSM2518777; mel_rep2: GSM2518778; mel_rep3: GSM2518779; mel_rep4: GSM2518780; mel_rep5: GSM2518781; mel_rep6: GSM2518782; mel_rep7: GSM2518783; Mouse: mm_rep1: GSM2518784; mm_rep2: GSM2518785.

### Single-cell RNA-seq: data processing, alignment and gene quantification

We chose read 1 to be 20 bp long to avoid reading into the poly(A) tail, leaving 64bp for read 2. The sequencing quality was assessed by FastQC (version 0.11.2). The base qualities of read 1 were particularly inspected, since they contain the cell and molecular barcodes and their accuracy is critical for the subsequent analysis. The last base of read 1 consistently showed an increase in T content, possibly indicating errors during bead synthesis. We observed a similar trend when re-analyzing the original data from Macosko et al. [7]. Part of these errors were handled and corrected as described later. For read 2, we used the Drop-seq tools v. 1.12 [7] to tag the sequences with their corresponding cell and molecular barcodes, to trim poly(A) stretches and potential SMART adapter contaminants and to filter out barcodes with low quality bases.

For mixed species experiments with human and mouse cells, the reads were then aligned to a combined FASTA file of the hg38 and mm10 reference genomes, using STAR [14] v. 2.4.0j with default parameters. Typically, around 65% of the reads were found to uniquely map to either of the species genomes. The *Drosophila melanogaster* sequences were mapped to the BDGP6 reference genome, typically around 85% and of the reads mapped uniquely. For the mouse hindbrain samples 75% of all sequence reads mapped uniquely. Non-uniquely mapped reads were discarded.

The Drop-seq toolkit [7] was further exploited to add gene annotation tags to the aligned reads (annotation used was Ensembl release 84) and to identify and correct some of the bead synthesis errors described above. The number of cells (cell barcodes associated with single-cell transcriptomes) was determined by extracting the number of reads per cell, then plotting the cumulative distribution of reads against the cell barcodes ordered by descending number of reads and selecting the inflection point (“knee”) of the distribution. It was similar to the number of single-cell transcriptomes expected during the Drop-seq run (see Suppl. Figure S1 for details). Finally, the DigitalExpression tool [7] was used to obtain the digital gene expression matrix (DGE) for each sample.

### Exploratory analysis, visualization and filtering of Drop-seq data

We developed an R software package (‘dropbead’; available for download at https://github.com/rajewsky-lab/dropbead, including a tutorial), which offers an easy and systematic framework for exploratory data analysis and visualization. Dropbead provides a function for computationally determining the inflection point and hence the number of cells in a sample. The starting point for subsequent analysis is the sample’s DGE. Dropbead provides functions for creating species separation plots and violin plots of genes and transcripts per cell. Dropbead can be used to easily filter and remove genes with low counts, cells with few UMIs, or to keep the best cells according to a certain criterion. Drop-bead was used to generate Figures 2a, 2b, 2c, 3a, 4a, S1a, S1b, S1f, S2a, S2b, S2c, S3a, S3b, S5a, S5b.

**Fig. 1.**
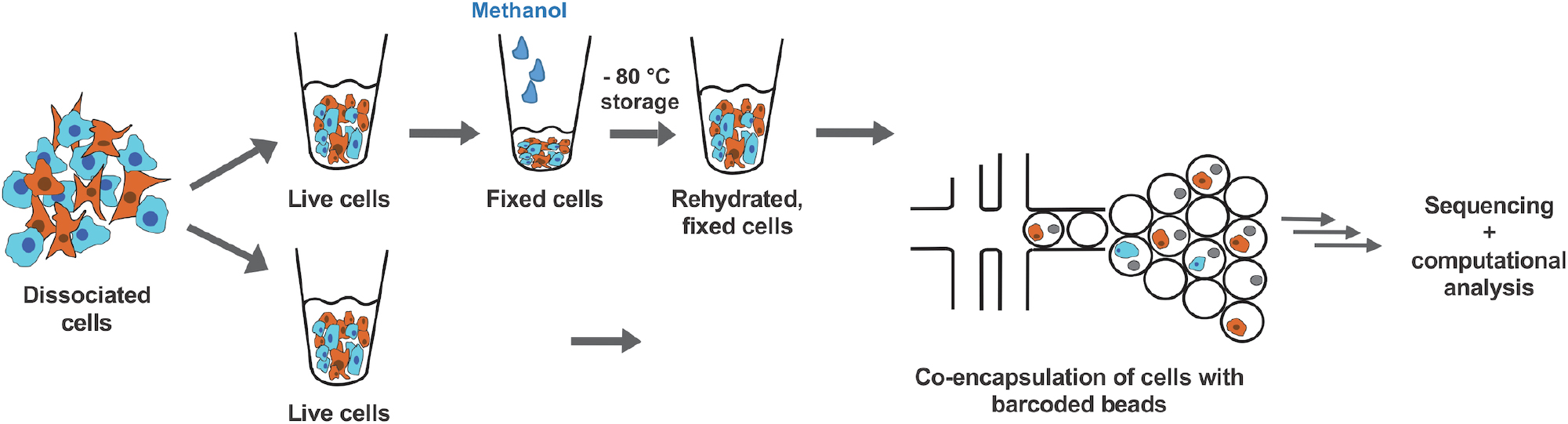
Cell preparation for droplet-based single-cell transcriptional profiling. Schematic of experimental workflow. Cultured human (HEK) and mouse (3T3) cells were dissociated, mixed and further processed to analyze the transcriptomes of either live or fixed cells by Drop-seq. Washed cells were gently resuspended in 2 volumes of ice-cold PBS, then fixed by addition of 8 volumes of ice-cold methanol. Methanol-fixed cells could be stored for up to several weeks at -80 °C. Prior to Drop-seq, cells were washed before passing them through a 35 − 40 μm cell strainer. Cells were then separately encapsulated in droplets together with a single bead in a microfluidic co-flow device and single-cell transcriptomes sequenced in a highly parallel manner. Downstream analysis and systematic quantitative comparisons were subsequently made from separate experiments using live or fixed cellular input material with an R package (‘dropbead’) that we developed and is freely available for download at https://github.com/rajewsky-lab/dropbead.

**Fig. 2.**
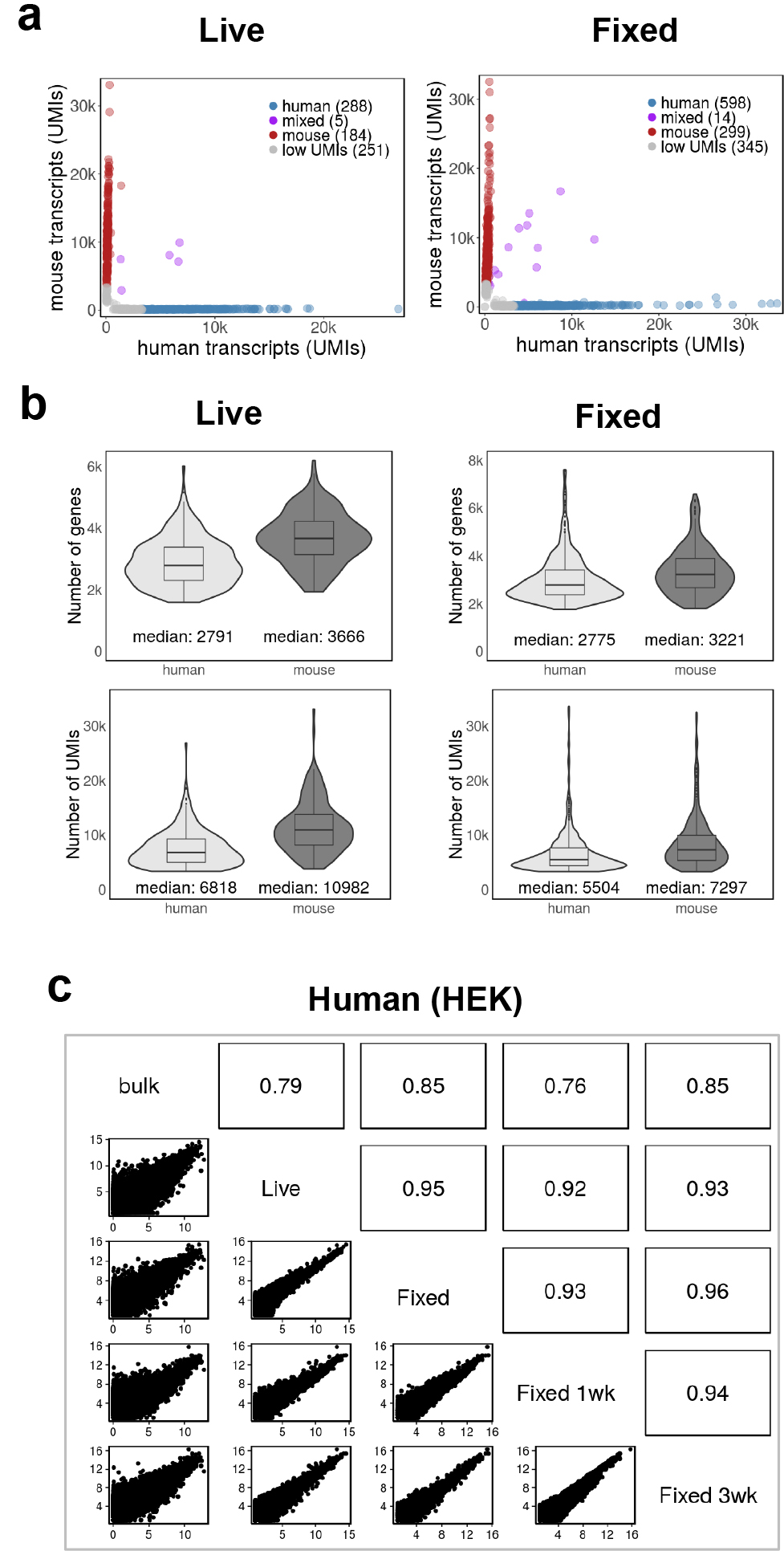
Transcriptome integrities and gene expression levels are preserved in fixed cells. **a** Drop-seq of mixed human and mouse cells (50 cells/µl). Plots show the number of human and mouse transcripts (UMIs) associating with a cell (dot) identified as human- or mouse-specific (blue or red, respectively). Cells expressing less than 3500 UMIs are grey, doublets are violet. **b** Distribution and the median of the number of genes and transcripts (UMIs) detected per cell (> 3500 UMIs). Libraries were sequenced to a median depth of ~20 500 (Live) and ~15 500 (Fixed) aligned reads per cell. **c** Gene expression levels from live and fixed cells correlate well. Pairwise correlations between bulk mRNAseq libraries (left) and Drop-seq single-cell experiments (right). Non-single cell bulk mRNA-seq data were expressed as reads per kilobase per million (RPKM). Drop-seq expression counts were converted to average transcripts per million (ATPM) and plotted as log2 (ATPM + 1). Upper right panels show Pearson correlation. The overlap (common set) between all 5 samples is high (17 326 genes). Experiments with live and fixed cells were independently repeated with similar results (unpublished).

**Fig. 3.**
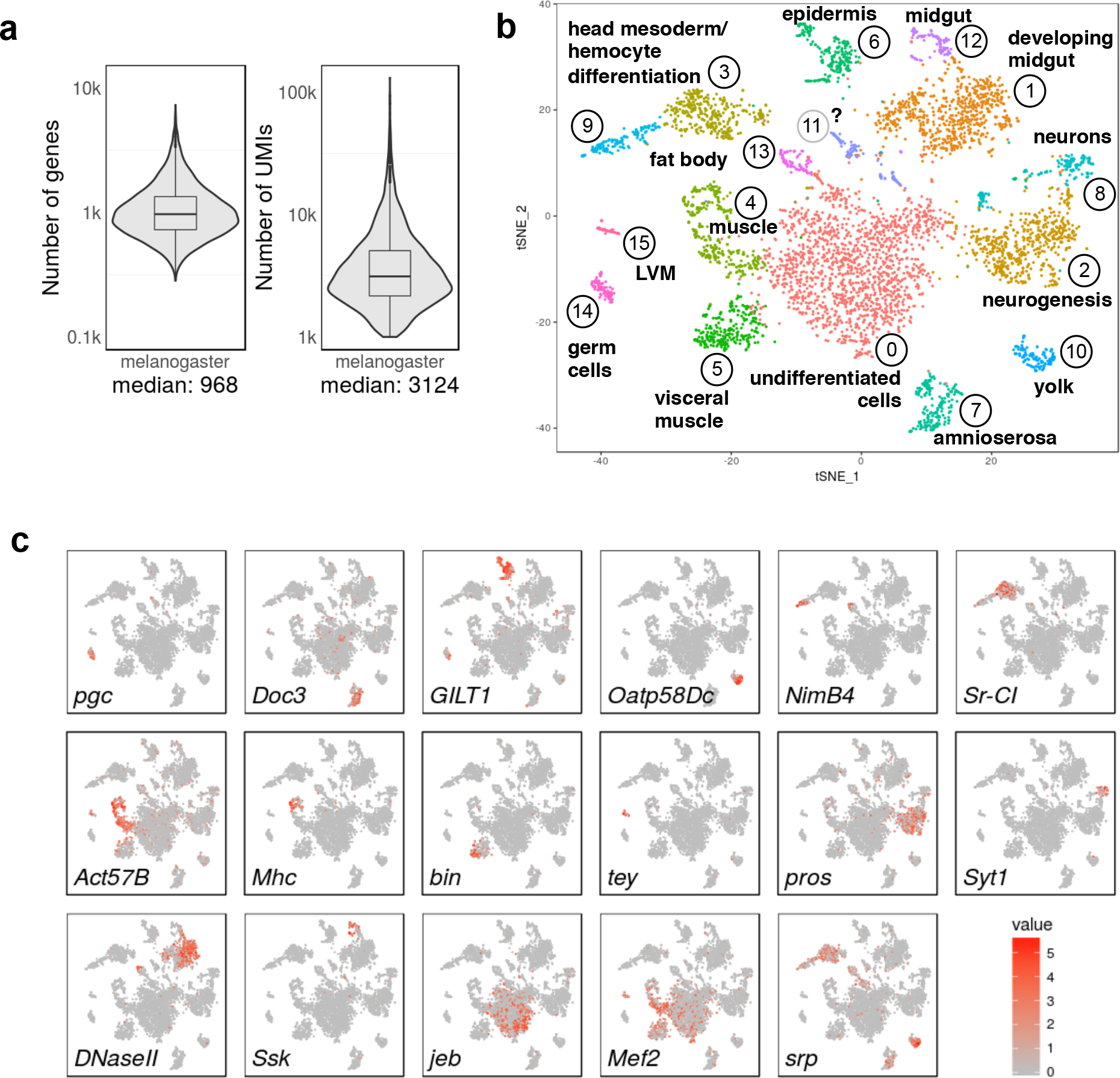
Primary, fixed cells from dissociated *Drosophila* embryos cluster into distinct cell populations. *Drosophila* embryos were collected in 2 hour time periods, aged for 6 hours, dissociated into single cells and fixed. Drop-seq data correspond to two independent embryo collections, with 3 and 4 technical replicates, respectively. Libraries were sequenced to an median depth of ~13 250 aligned reads per cell. Cells expressing less than 1 000 UMIs were excluded from the analysis. **a** Distribution and the median of the number of genes and transcripts (UMIs) detected per cell in Drop-seq data pooled from seven Drop-seq runs, representing a total of 4 873 cells. Note that violin plots are displayed on a log scale. **b** Clustering of 4 873 fixed cells into distinct cell populations. The plot shows a two-dimensional representation (tSNE) of global gene expression relationships among all cells. Tissue associations were made by ImaGO term analysis [19] on the 50 most variable genes of each cluster (Table S1), followed by inspection of publicly accessible RNA *in situ* staining patterns. LVM, longitudinal visceral musculature. **c** Marker gene expression in clusters of *Drosophila* embryo cells (see text for explanations). Expression colored based on normalized expression levels.

**Fig. 4.**
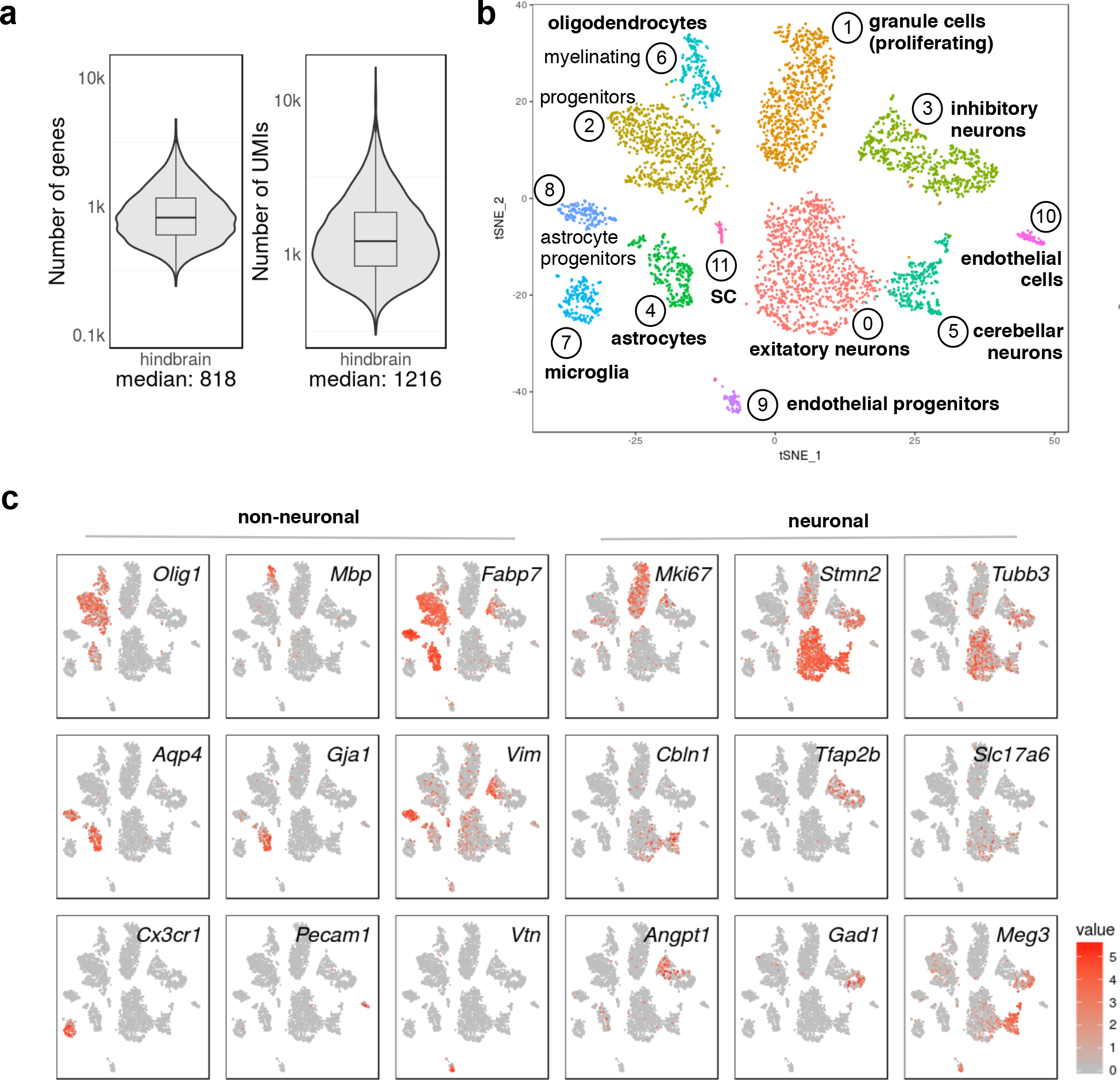
Sorted, fixed mouse brain cells allow identification of distinct neural and non-neural cell types. Hindbrains and cerebellum from newborn mice were microdissected and dissociated, and cells were sorted by FACS into methanol and stored. Drop-seq data correspond to two independent biological replicates. Libraries were sequenced to a median depth of ~7 100 reads per cell. **a** Distribution and the median of the number of genes detected per cell (> 300 UMIs) in Drop-seq data pooled from two Drop-seq runs, representing 4 366 cells. Note that violin plots are displayed on a log scale. **b** Clustering of 4 366 fixed cells into distinct cell populations marked by color (Table S2). The plot shows a two-dimensional representation (tSNE) of global gene expression relationships among all cells. Tissue associations of cell clusters were identified by assessing the 50 most variable genes in each cluster and confirmed by inspection of publicly accessible images of RNA in situ hybridizations. **c** Known marker gene expression in clusters of brain cells (see text for explanation). Expression colored based on normalized expression levels.

We discarded cells from subsequent analysis, which had less than 3 500 UMIs (HEK and 3T3 cells), 1 000 UMIs (*Drosophila* samples), or 300 UMIs (mouse samples). In the human-mouse mixed species experiments, a threshold of 90% (90 out of 100 UMIs for one species) was selected to confidently declare a cell as being either of the species and not a human/mouse doublet. In order to assess whether fixation generates “low-quality cells” [15], we determined the proportion of non-mitochondrial reads: for every cell we computed the sum of UMIs corresponding to RNA encoded by the mitochondrial genome and then subtracted this number from the sum of all UMIs in that cell. We divided this number by the total number of UMIs in the cell to obtain the non-mitochondrial content as a percentage for every sample.

### Single-cell RNA-seq: filtering, normalization and correlations of gene expression levels

The raw counts in the DGE were normalized to average transcripts per million (ATPM) as follows: the UMI counts for every gene in a given cell were divided by the sum of all UMIs in that cell. These counts were then multiplied by the sum of all UMIs of the cell with the highest number of UMIs in that library. Correlations of gene expression levels between single-cell samples were computed by first subsetting the DGEs of the two samples to the intersection of the genes captured in both libraries and then computing the sum of gene counts across all cells in each library. Plotting of correlations is shown in log-space. For the correlation of Drop-seq data against mRNA-seq, we converted gene counts into RPKMs and used the mean value of all isoforms lengths for a given gene. For all corrrelations, the intersection (common set) of genes was high, around 17 000 genes for human and mouse samples (cell lines and primary cells), and 10 000 genes for *D. melanogaster*.

### Bulk mRNA-seq libraries

Live, cultured cells (Flp-In T-Rex 293 HEK cells, NIH 3T3 cells), intact, live *Drosophila melanogaster* embryos, and sorted, methanol-fixed cells from dissected newborn mouse hindbrain and cerebellum were used for total RNA extraction with TRIZOL. Strand-specific cDNA libraries were generated according to the Illumina TruSeq protocol (TruSeq Stranded mRNA LT Sample Prep Kit, Illumina, San Diego, USA) using between 24 to 260 ng of total RNA input. The 1.8 pM libraries were sequenced on an Illumina NextSeq 500 System, using High Output v2 Kit (150 cycles), single read 150bp, index read: 6 bp.

Unique identifiers: bulk_hek: GSM2518786; bulk_3t3: GSM2518787; bulk_mel1: GSM2518788; bulk_mel2: GSM2518789; bulk_mm: GSM2518790.

### Single-cell RNA-seq: clustering and identification of cell populations and marker genes

For the *Drosophila* embryos and mouse hindbrain samples, after filtering our samples with dropbead we used Seurat [16] for cluster analysis. We first identified a set of highly variable genes, which we used to perform principal component analysis (PCA). Judged by their statistical significance, the first few (~30-50) PCs were subsequently used as inputs for clustering and subsequent t-distributed stochastic neighbour embedding (tSNE) representation. We used Seurat to identify the marker genes for each of the clusters in the tSNE representation.

The initial clusterings for both the *Drosophila* embryos and the mouse hindbrain samples contained a small number of cell clusters which were difficult to characterize (three and one cluster, respectively). After close inspection we identified two of them in the *Drosophila* case as being nuclei, as those cells were lacking substantial expression of mitochondrially encoded genes compared to the rest of the cells. We classified these cells as nuclei (generated by mechanic disruption in the cell isolation procedure) and excluded them from further analysis.

Furthermore, extrapolating from our mixed species experiments with human and mouse cell lines, we anticipated around 10% of same- species doublets for the combined *Drosophila* embryos and mouse hindbrain samples. In the first case, we identified a set of cells which co-expressed marker genes known to be expressed exclusively in separate tissues. In the latter case, we identified a similar set of cells which additionally contained higher portions of ribosomal protein coding mRNAs than the rest of the cells. We reasoned that both sets were likely cell doublets and excluded them in order to perform the final cluster analysis shown in Fig. 3b and Fig. 4b.

## RESULTS

### Methanol fixation preserves single-cell transcriptomes for droplet-based sequencing

#### Drop-seq with methanol-fixed cells allows correct species assignments in species-mixing experiments

In order to assess whether methanol fixation is compatible with Drop-seq, we adapted our previously developed methanol fixation protocol [12] to adherent, mammalian cell lines (see Materials and Methods for details). Methanol-fixed cells remained visible under the microscope as single, intact round cells, which disappeared upon addition of Drop-seq lysis buffer due to complete cellular lysis (data not shown). Fixation did not induce a microcopically detectable increase in cell doublets.

To assess the quality of single-cell transcriptomes generated by the Drop-seq procedure after methanol fixation, we used a mixture of cultured human (HEK) and mouse (3T3) cells as performed previously [7]. Both, live and fixed cell mixtures were used at a final concentration of 50 cells/µl for Drop-seq runs carried out on the same day, and cDNA libraries were processed in parallel. Figure 1 shows the experimental workflow and Figure 2 the results of this experiment. We counted the numbers of human and mouse transcripts (UMIs) that were associated with each cell barcode. Both, live and fixed cells, could be confidently assigned to their species of origin using a threshold of 90% species-specific transcripts (Fig. 2a), suggesting that methanol fixation preserves cell integrity and the species-specificity of a cell’s transcriptome. In addition, this experiment showed that fixation did not substantially increase human/mouse cell doublet rate.

In Drop-seq, cell numbers are selected computationally from the inflection point (“knee”) in a cumulative distribution of reads plotted against the cell barcodes ordered by descending number of reads. Cell barcodes beyond the inflection point are believed to represent “ambient RNA” (e. g. contaminating RNA from damaged cells), not cellular transcriptomes [7]. As shown in Suppl. Figure S1a, our fixation protocol did not interfere with our ability to computationally select cells.

#### Transcript and gene numbers from live and fixed cells are similar

Figure 2b shows the number and distribution of genes and transcripts (UMIs) in live and fixed cells. Median transcript and gene numbers from fixed cells and their distributions were similar to live cells, indicating that methanol fixation did not change the sensitivity of Drop-seq results (Fig. 2b, Suppl. Fig. S1b). Processing and sequencing a lower number of transcriptomes from the same Drop-seq experiments resulted in higher gene and transcript numbers in both cases (Suppl. Fig. S1b).

#### Gene expression levels correlate well between live and fixed cells

We treated single-cell transcriptomes as a bulk population and plotted transcript counts from fixed cells against transcript counts from live cells, to determine how well they correlate. Figures 2c and S2c show that gene expression levels from live and fixed cells were highly correlated (R ≥ 0.95). Furthermore, transcripts from live and fixed cells against human (HEK) and mouse (3T3) cell bulk mRNA-seq data correlated well (R ≥ 0.79; Fig. 2c and Suppl. Fig. S2c).

Taken together, our data suggest that methanol fixation faithfully preserves single-cell transcriptomes for the Drop-seq procedure.

### Fixed cells can be stored for weeks to give reproducible Drop-seq results

We tested whether fixed cells can be stored, and if so, for how long they can be used for Drop-seq experiments. In order to address this question, we fixed cells and stored them at -80°C for one week or 3 weeks. As shown in Suppl. Figures S1 and S2, single-cell transcriptomes from cells stored for either 1 or 3 weeks performed well in experiments with mixed human and mouse cells. Our results were robust with respect to computational cell selection (Suppl. Fig. S1a), the ability to assign barcodes to an individual cells’ organism of origin (Suppl. Fig. S2a), and the median number of genes and transcripts per cell (Suppl. Fig. S2b).

Finally, gene expression profiles from fixed cells stored for 1 or 3 weeks correlated well with each other and those of cells that were fixed immediately prior to Drop-seq (Fig. 2c and Suppl. Fig. S2c). They also showed high correlation with bulk mRNA-seq data (Figure 2c and Suppl. Fig. S2c). We concluded from these data, that fixed cells are stable for several weeks and can be used for Drop-seq experiments without loss in quality.

### Methanol fixation preserves RNA integrity and cytoplasmic RNA content

#### High-quality RNA and cDNA libraries can be prepared from fixed cells

Suppl. Figure S1c shows that high-quality, intact total RNA could be extracted from fixed cells after storage in 80% methanol for 20 weeks (RNA was extracted from the same batch of fixed cells that were used to generate the results shown in Suppl. Fig. S2). Furthermore, Suppl. Figure S1 shows BioAnalyzer traces corresponding to all four cDNA libraries analyzed in this study: cDNA libraries from methanol-fixed cells appeared indistinguishable from cDNA libraries obtained from live cells (Suppl. Fig. S1d and unpublished data). Additionally, we confirmed that cDNA libraries from fixed cells did not contain a major ‘hidden’ peak of low molecular weight fragments normally removed by the library clean-up procedure (Suppl. Fig. S1e and unpublished data).

#### Mitochondrially-encoded transcripts are not elevated in methanol-fixed cells

An increase in the proportion of transcripts from mitochondrial genes (37 mtDNA-encoded mRNAs) is believed to indicate low quality cells that are broken or damaged to varying degrees [15]. It is thought that this phenomenon is caused by leakage leading to relative loss of cytoplasmic mRNAs compared to mitochondrially located mRNA transcripts, which are protected by two mitochondrial membranes. As shown in Suppl. Figure S1f, we observed less than 10% loss of cellular cytoplasmic mRNA content across all three fixed Drop-seq libraries. Thus, fixation does not seem to cause a major increase in “low quality cells”.

Taken together, our data indicate that methanol fixation is able to preserve RNA integrity, and subsequent cDNA library generation during the Drop-seq procedure. Our results also show that fixed cells can be stored for prolonged periods up to at least several weeks or months.

### Methanol fixation allows cell type identification in developing *Drosophila* embryos

#### Fixed, primary low RNA content cells from dissociated Drosophila embryos perform well in Drop-seq

Primary cells tend to be smaller and contain less RNA than cultured cells, making them harder to analyze by single-cell sequencing [17]. Therefore, we first tested how methanol fixation and subsequent storage performs on primary Drosophila cells, which are generally much smaller than most mammalian cell types [18]. Figure 3 and Suppl. Figure S3 show the results from seven Drop-seq runs (3 and 4 technical replicates, respectively) performed with two independently collected and processed samples from *Drosophila* embryo pools collected over a 2-hour period of time and aged to assure a rich mixture of differentiating and differentiated cell types (about 75% of developmental stages 10 and 11). The resulting single-cell sequencing data allowed computational selection of cells from “knee plots” (Fig. S3a), and cross-correlations of aggregated reads were highly reproducible (R≥0.96 across the 7 replicates and R≥0.82 for comparisons with bulk mRNA-seq samples; Fig. S3b and S4b). At a sequencing depth of a median of ~13 250 aligned reads per cell (filtering cells with >1000 UMIS), we obtained a median of ~1000 genes and ~3000 transcripts (UMIs) per cell (Fig. 3a), indicating that fixation is suitable for primary cells with low RNA content.

#### Methanol fixation allows cell type identification in developing Drosophila embryos

After removing nuclei (characterized by underrepresentation of mitochondrially encoded genes) and likely cell doublets (see Methods), we performed PCA and 2D clustering by tSNE using the remaining 4 873 cells (Figure 3b and Suppl. Fig. S4). Variance was captured in many principal components across distinct embryonic cell populations (Fig. S4a). Clustering analysis revealed numerous cell clusters most of which could be associated with developing tissues and cell types, based on gene expression profiles (Fig. 3b, Table S1). Both samples and all Drop-seq runs contributed to the observed clusters, indicating high reproducibility between biological and technical replicates (Fig. S4b).

Tissues were assigned through imaging gene ontology (ImaGO) term analysis [19] of the 50 most variable genes in each cluster (Table S1) as a first approximation, followed by inspection of publicly available RNA *in situ* staining patterns [20] of highly variable, as well as other known tissue-specific genes. We identified one cluster encompassing an assemblage of undifferentiated cells (cluster 0, marked by genes such as *jelly belly, jeb*). Other clusters comprised cell identities corresponding to the germ line and derived from all 3 germ layers. Known cell-type markers show cluster specific expression patterns as expected (Fig. 3c) such as in germ cells (*polar granule component, pgc*), amnioserosa (T-box transcription factor Dorsocross3, *Doc3*), epidermis (disulphide oxyreductase, *GILT1*), and yolk (*Oatp58Dc*). Clusters 3, 4, 5, 9, 13 and 15 all comprise mesodermally derived cells, albeit in distinct spatial and developmental sub-populations: Cluster 3, 9 and 13 appear to constitute subpopulations of the fat body (pathogen receptor *NimB4*) and head mesoderm giving rise to differentiating hemocytes/macrophages (scavenger receptor Class C type I, *Sr-CI)*. Clusters 4 and 5 include somatic and visceral musculature, but appear to separate the differentiation state of the developing muscle: more differentiated cells in cluster 4 expressing contractile machinery (myosin heavy chain, *Mhc*), whereas cluster 5 cells appear less differentiated (*binou, bin*). Cluster 15 seems to specifically identify the cells of longitudinal visceral musculature (*tey*). Similarly, ectoderm clusters 2 (*prospero, pros*) and 8 (Synaptotagmin 1*, Syt1*) both comprise neurogenic cells, but cluster 2 cells appear to be developmentally less differentiated. Accordingly, biological process GO term enrichment for cluster 2 terms were more generally linked to nervous system development, whereas cluster 8 was linked to synaptic signalling and other terms indicating a functioning nervous system. Clusters 1 (*DNAse II*) and 12 (Snakeskin*, Ssk*) are both endodermal cell populations, but while cluster 1 primarily constitutes the mid- and hindgut primordium, cluster 12 contains more differentiated, functional cells of the gut. Lastly, *myocyte enhancing factor* (*Mef2*) and *serpent* (*srp*) are two examples of transcription factors, which are expressed in distinct mesoderm-derived clusters, as expected at this stage of development.

### Sorted, fixed mouse brain cells allow identification of distinct neural and non-neural cell types

In order to address the question whether methanol fixation can be used in conjunction with FACS, we dissected hindbrain and cerebellum from newborn mice, dissociated the cells and sorted live, propidium iodide-negative cells directly into ice-cold methanol. We expected to obtain a mixture of neurons, glia and non-neuronal cell types for further analysis. Two independent biological replicates allowed computational cell selection from “knee plots” (Fig. S5a, b) and were highly reproducible (R=0.95; R≥0.8 for comparisons with independently prepared bulk mRNA-seq data). After combining the data and removing low quality cells (expressing <300 UMIs) as well as cell doublets, we obtained a median of ~800 genes and ~1200 transcripts (UMIs) per cell for the remaining 4 873 cells, even though the samples were sequenced only at a shallow depth (a median of ~7 100 aligned reads per cell; Fig. 4a). PCA revealed variance captured in many principal components (Fig. S6a) and 2D representation by tSNE produced 12 clusters all of which contained readily identifiable cell types (Fig. 4b; Table S2). Both biological replicates contributed to all observed clusters (Fig. S6b).

Cell populations were identified through their gene expression signatures (Fig. 4c; Table S2) and encompassed neurons and glial cells. We identified different neuronal cell types such as proliferating granule cells (proliferation marker *Mki67*; neuronal marker stahmin-like 2, *Stmn2*), excitatory neurons (glutamatergic neuronal marker *Slc17a6*/*Vglut2* and neuronal markers *Stmn2* and tubulin beta 3, *Tubb3*), inhibitory neurons (GABAergic neuronal marker *Gad1*/*Vgat1*; *Tubb3*; transcription factor AP2 beta, *Tfap2b*) and cerebellar neurons (Cerebellin, *Cbln1*; lncRNA *Meg3*; *Stmn2*). Among the glial cells, we identified oligodendrocyte progenitors (oligodendrocyte transcription factor 1, *Olig1*; fatty acid binding protein 7, *Fabp7*), myelinating oligodendrocytes (myelin basic protein, *Mbp*; *Olig1*), microglia (chemokine receptor Cx3cr1), and astrocytes (gap junction protein alpha 1, *Gja1*; aquaporin 4, *Aqp4*; *Fabp7*) as well as astrocyte/neuronal progenitors (vimentin, *Vim*; *Aqp4, Fabp7*). We also identified a subtype of myelinating glia (cluster 11), which expressed myelin protein zero (*Mpz*), probably Schwann cells from cranial nerves entering the hindbrain (Fig. S6c). In addition, we found non-neural cell types such as endothelial cells expressing platelet/endothelial adhesion molecule 1 (*Pecam1*), and endothelial progenitors (vitronection, *Vtn*). Markers were confirmed to be expressed in newborn hindbrain and cerebellum by inspecting RNA *in situ* hybridization images in publicly available databases (The Gene Expression Database (GXD) [21], available at www.informatics.jax.org/expression.shtml; © 2008 Allen Institute for Brain Science. Allen Developing Mouse Brain Atlas. Available from: http://developingmouse.brain-map.org/)

Together, our data demonstrate that methanol can be used to fix and store primary cells for Droplet-based sequencing, including low input RNA cells such as differentiating embryonic *Drosophila* cells and a wide variety of mammalian brain cells, including neuronal subtypes.

## DISCUSSION

Few studies so far have explicitly dealt with cell preservation protocols for single-cell sequencing. One study uses cryopreservation followed by flow cytometry to sort single cells for subsequent processing [22]. While cryopreservation is compatible with droplet-sequencing in principle [17], it remains to be determined how well Drop-seq will perform with recently thawed cells that may be fragile and prone to die. Another study describes fixation of cells by cross-linking with formaldehyde [23], followed by reverse cross-linking (breakage of methylene bridges between protein and RNA molecules with heat). Crosslinking induces chemical modifications that inhibit poly(dT) annealing, reverse transcriptase and cDNA synthesis making cross-link reversal necessary [10]. However, reversal of crosslinking is often inefficient, leaves adducts and can be expected to result in high loss of available RNA molecules in a non-uniform manner.

We have shown here that a simple methanol fixation of tissue culture cells does not lead to significant RNA loss or degradation, and is easily compatible with the established Drop-seq single-cell sequencing protocol and workflow. It is possible to store single-cell suspensions for prolonged times at low temperatures and, therefore, to separate the sample preparation phase in time or location from the actual droplet-sequencing procedure. In addition to the cell culture lines used in this study (human HEK and Hela cells, mouse NIH3T3 cells), we have successfully applied our methanol fixation protocol to a variety of other cell lines or cultured cells (HeLa cells, mouse cycling and non-cycling pre-B and primary mouse lymphoma cells).

Beyond cultured cells, which in many instances contain more RNA than oftentimes smaller, primary cells [17], Drop-seq can be successfully performed with fixed cells from complex tissues such as dissociated, later stage *Drosophila* embryos and mouse brain, as shown in this study. Single-cell sequencing data from individual runs of primary tissues from both organisms yielded highly reproducible results. Even though we applied only a relatively shallow sequencing depth (a median of ~7 100 aligned reads per cell) to our mouse brain sample, the single-cell sequencing data allowed identification of diverse cell types and subpopulations of cells, including subcluster differentiation and developmental trajectories (Fig. 4c) [24, 25]. For example, a large cluster of inhibitory neurons (marked by *Gad1, Tfap2b*; cluster 3 in Fig. 4b, c) contained less mature, still dividing cells in the left part (marked by *Vim, Mki67* and *Angpt1*) and more mature neurons in the right part (*Tubb3, Stmn2, Meg3*). Furthermore, in the oligodendrocyte clusters (2 and 6 in Fig. 3b) more mature, myelinating oligodendrocytes clustered to the upper left of cluster 6 (marked by *Mbp, Olig1*), newly formed oligodendrocytes in the middle of cluster 2 (*Olig1, Fabp7*) and still dividing progenitors to the lower right of cluster 2 (marked by *Mki67, Fabp7, Olig1*).

We also found that single-cell sequencing data from methanol-fixed cells were of sufficient quality to carry out spatial mapping and 3D reconstruction of a virtual *Drosophila* embryo at the onset of gastrulation [13]. Methanol fixation allowed us to prepare and store cells from carefully staged, early embryos incrementally in small batches.

However, methanol fixation may not work in all circumstances, for all tissues or cell types. Successful fixation may be challenging especially for tissues with a high content in proteases and RNAses such as pancreas, gall bladder, skin or lymphatic and immune tissues. In support of this notion, we observed a failure to generate Drop-seq cDNA libraries and to isolate intact RNA from fixed cells of a mouse lymphoma *ex vivo* (unpublished results). For these types of tissues, it will be important to determine at which step RNA degradation occurs, before or after fixation. Modifications to the fixation protocol such as addition of RNAse inhibitor during the rehydration step (as used in our experiments with primary cells) may remedy these problems. It also remains to be determined whether methanol-fixation is compatible with the ‘InDrop’ protocol [8], another, recently developed droplet-based sequencing approach that involves a different detergent for cell lysis and cDNA library construction. A recent study that uses combinatorial indexing to transcriptionally profile large numbers of single cells relies on methanol fixation [26], suggesting that alcohol-based fixation may be compatible with a wider range of single-cell sequencing approaches.

## CONCLUSIONS

The availability of a simple cell fixation protocol will open up many previously inaccessible experimental avenues for droplet-based single-cell transcriptomics. Fixation and preservation of cells at an early stage of preparation removes bias and technical variation, prevents cell stress, or unintended ageing during the experiment, and facilitates systematic assessment of experimental parameters. Cell fixation may also significantly ease the logistic coordination of large-scale experiments. In a variety of situations, fixation may be the only solution to being able to process and provide cellular input material: Examples are rare cells which cannot be obtained in one experimental session [13], clinical specimen which require transportation, or cells that are hard to isolate, and require extensive upstream processing such as tissue dissociation followed by flow cytometry. In summary, we expect that the methanol-based cell fixation procedure presented here will greatly stimulate high-throughput single-cell sequencing studies in diverse areas.

## Declarations

### Ethics approval

Mouse breeding, housing and experiments were conducted in accordance with institutional German regulations under permits for C.B.

### Consent for publication

Not applicable.

### Availability of data and materials

The data sets supporting the conclusions of this article are available in the GEO repository (record GSE89164).

Software is available at https://github.com/rajewsky-lab/dropbead

## Acknowledgements

We thank Rahul Satija for advice with single-cell sequence analysis, Evan Macosko, Melissa Goldman and Steve McCarroll for help with setting up Drop-seq, Sara Formichetti for RNA extractions, Haiyue Liu for testing our R scripts, and our colleagues from the Rajewsky laboratory for manifold help and critical discussions. We also thank Petra Stallerwo for the mouse husbandry, and Hans-Peter Rahn for help with sorting of mouse hindbrain cells.

## Author’s contributions

JA, CK, NR conceived the study; CK, NR defined the broader strategy and supervised; JA, SP, NK, SG, PW, PLR, CB, RZ, CK, NR designed the experimental and analytical strategies; NK developed computational tools, performed all the computational analyses; PW, PLR, SA, LS, AB prepared biological materials; JA, SP, SG, SA, LS, AB, CK, NR performed single-cell sequencing and library preparations; JA, SP, NK, SG, PW, PLR, CB, RZ, CK, NR analyzed data; CB, RZ, NR procured funding; JA, SP, NK, PW, PLR,CB, RZ, CK, NR wrote, discussed and edited the manuscript and prepared the figures. All authors read and approved the final manuscript.

## Author details

Not applicable.

## ADDITIONAL FILES

**Suppl. Fig. S1 (related to Fig. 2)**

**Suppl. Fig. S2 (related to Fig. 2)**

**Suppl. Fig. S3 (related to Fig. 3)**

**Suppl. Fig. S4 (related to Fig. 3)**

**Suppl. Fig. S5 (related to Fig. 4)**

**Suppl. Fig. S6 (related to Fig. 4)**

**Suppl. Table S1** Top 50 marker genes expressed in 4 873 fixed, primary cells from *Drosophila* embryos

Suppl. Tables S1 and S2 contain the top 50 marker genes per cluster, provided by Seurat’s function findAllMarkers [16]. We additionally ordered them per cluster in decreasing log2-fold change (log2FC). The Log2FC was computed for a given gene by dividing its average normalized expression for a given cluster over the average normalized expression in the rest of the clusters and taking the logarithm of the fold change.

**Suppl. Table S2** Top 50 marker genes expressed in 4 366 sorted, fixed cells from mouse hindbrain and cerebellum

